# Verified the effectiveness of AsCpf1 system in a variety of vertebrate species

**DOI:** 10.1101/272716

**Authors:** Zhuo Li, Zhaoying Shi, Nana Fan, Yongqiang Chen, Jing Guo, Jingchun Wu, Hong Song, Shilong Chu, Kunlun Mo, Bentian Zhao, Zhen Ouyang, Dandan Tian, Shaoyang Zhao, Jieying Zhu, Jiekai Chen, Yonglong Chen, Liangxue Lai, Duanqing Pei

## Abstract

CRISPR/Cpf1 system is a novel genomic editing tool. Because of its more sophisticated components, and lower off-target rate, it has the potential to become a better gene-editing tool. Previous reports showed that CRISPR/Cpf1 could work effectively in multiple species. But our data show that AsCpf1 activity has a big difference in different vertebrates. Using in vitro experiments, we finally learned that the difference between species is due to temperature.

---

In recent years, gene-editing tools have been greatly developed, and are widely used. CRISPR/Cpf1 system is a new member of the gene-editing technique family, and current researches have shown its better ease of use(Jiang et al., 2017; Kim et al., 2016b; Port and Bullock, 2016; Tang et al., 2017; Zetsche et al., 2015). This system can target and cut specific double-stranded DNA (dsDNA) with only two elements: CRISPR RNA (crRNA) and Cpf1 protein. CRISPR RNA recognizes and targets specific DNA sequence, and guides Cpf1 protein to assemble a ternary complex; ultimately, Cpf1 protein cut the targeted dsDNA. Cpf1 protein is smaller than Cas9, and crRNA, only 43nt(including 23nt recognition sequence), is also shorter than sgRNA. Based on reports, Cpf1 has outstanding performance in the off-target profile(Kim et al., 2016a; Kleinstiver et al., 2016). These advantages have shown this system is potentially better than ever before gene-editing tools. From plants to animals, CRISPR/Cpf1 has been validated in multiple species(Kim et al., 2016b; Port and Bullock, 2016; Tang et al., 2017; Toth et al., 2016; Zetsche et al., 2015), but some commonly used laboratory animal models have been reported rarely. Thus, we want to test the CRISPR/Cpf1 system on embryos of four commonly used model animals: mice, porcine (female embryos), frogs(Xenopus), zebrafish; in this study, we have chosen AsCpf1, because of its significant functionality of cutting dsDNA(Kim et al., 2016a; Kim et al., 2016b; Kleinstiver et al., 2016; Zetsche et al., 2015), as testing subject on embryos of these four commonly used models.

In the mouse genome, we chose m*Tet2* locus as target gene loci, and selected four targets for this locus. Validated CRISPR/Cas9 target in m*Tet2* was used as positive control(Wang et al., 2013). In order to obtain more reliable results in mouse embryos, we first assessed the effectiveness of selected target sites in mouse embryonic stem cells (mESCs) (**Supplementary Figure 1a, b**). We transfected PiggyBac plasmids that contain AsCpf1 gene and puromycin resistance gene into mESCs. After puromycin selection, transgenic cells stably expressed AsCpf1 proteins. Then, the crRNAs were transfected into transgenic cells. 48 hours after transfection, cells were collected to do double strand DNA breaks (DSB) assay. The results show all m*Tet2* targets were cut (**Supplementary Figure 1c, d**), and the highest cleavage efficiency is crm*Tet2*-1, 50% (**Supplementary Figure 1c, Supplementary Table 1**); sgRNA cleavage efficiency at the same gene locus is 95% (**Supplementary Figure 1c, d**). After screening, we chose crm*Tet2*-1 as target for testing CRISPR/Cpf1 system in mouse embryos.

We optimized the nuclear localization signal (NLS) prior to the mouse experiment (**Supplementary Figure 1e**), and found that the addition of NLS at the N-terminus of AsCpf1 did not significantly alter the function of AsCpf1 (**Supplementary Figure 1f, g**). So in the follow-up experiments we only add NLS to the C-terminus.

Referring CRISPR/Cas9 mouse embryo operating standard(Wang et al., 2013), 50ng/ul AsCpf1 mRNA and 20ng/ul crRNA were used. Injected embryos were cultured to the blastocyst stage, and were collected to prepare genome. After Sanger sequencing, we found that crm*Tet2*-1 target site has been cut and had indel (efficiency: 32%) (**Fig a**, **Supplementary Figure 2a, Supplementary Table 2**).

**Figure.**
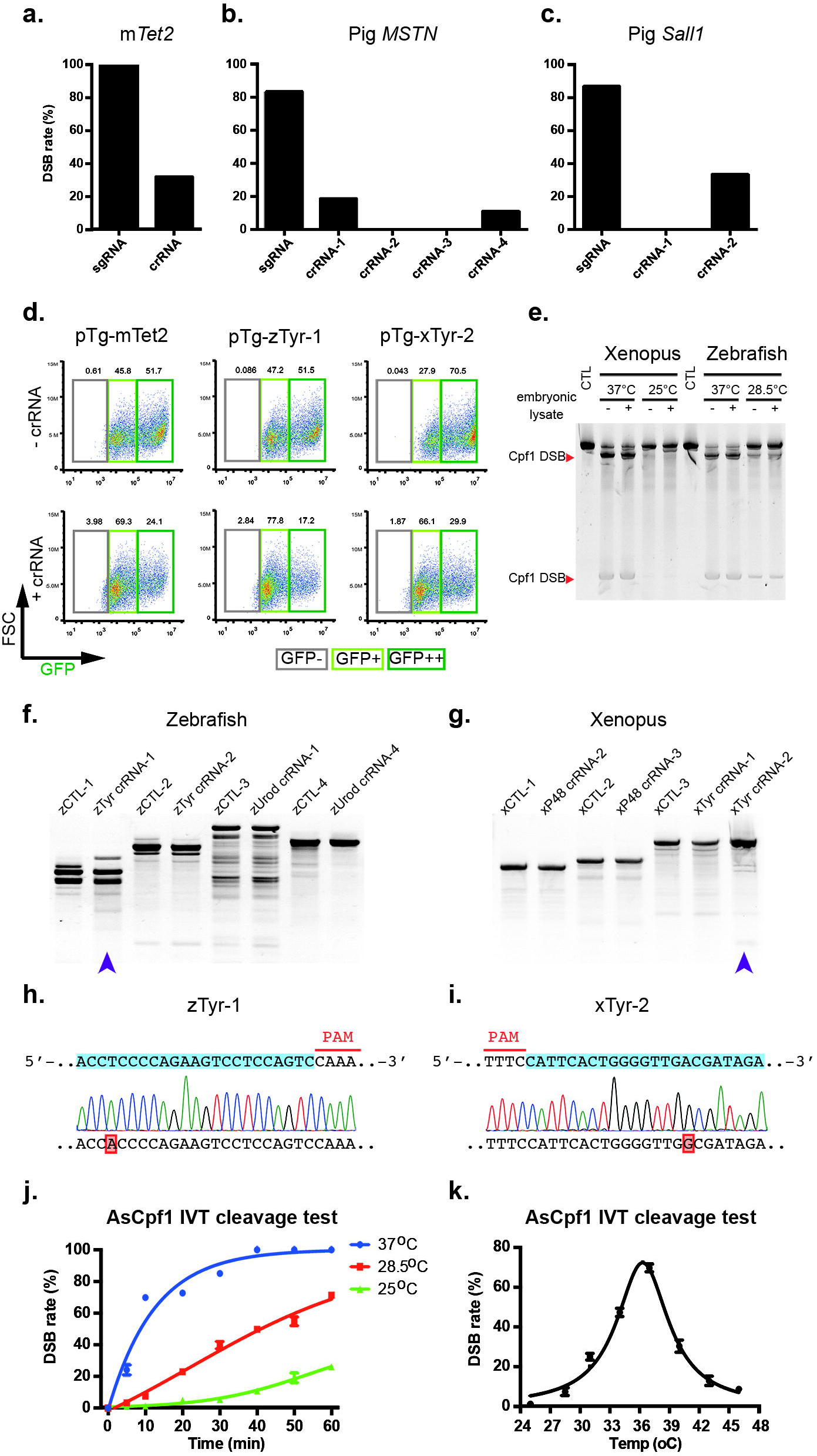
DNA cleavage efficiencies of AsCpf1 and SpCas9 targets at mouse *Tet2* locus in mouse embryos(**a**), at pig *MSTN* and *Sall1* loci in pig embryos (**b**), (**c**). (**d**) Detected mouse, zebrafish and xenopus exogenous targets by ETD system in HEK293T cells. (**e**) DNA cleavage efficiencies of linear pTg plasmids in zebrafish and xenopus embryo lysates mediated by AsCpf1 protein, AsCpf1 DNA cleavages are indicated by orange arrows. DNA cleavage efficiencies of AsCpf1 targets at zebrafish *Tyr* locus in zebrafish embryos(**f**), at xenopus *Tyr* locus in xenopus embryos (**g**) (suspected results are marked by purple arrow). Representative SNV caused by AsCpf1 in zebrafish (**h**) and xenopus Embryos (**i**). (**J**) AsCpf1 in vitro enzyme activity assay, DNA cleavage efficiencies have been checked at eight time points and there different temperatures. (**k**) AsCpf1 enzyme activity curve in the range of 25 °C to 46 °C, reacted in 10 minutes respectively.

In pigs, we have directly operated on parthenogenetic embryos. Four target sites at p*MSTN* locus and two at p*Sall1* locus have been chosen. Validated CRISPR/Cas9 targets in p*MSTN* and p*Sall1* were used as positive control. Referring CRISPR/Cas9 pig parthenogenetic embryo operating standard(Zhou et al., 2015), 200ng/ul Cpf1 mRNA and 20ng/ul crRNA were used. Two target sites of p*MSTN* have occurred DSB while their cleavage efficiencies are both lower than 20% (**Fig b**, **Supplementary Figure 2b, c, Supplementary Table 3**). Another target site at p*Sall1* locus also has DSB and it has a little higher cleavage efficiency - 33% (**Fig c**, **Supplementary Figure 2d, Supplementary Table 4**).

In zebrafish and xenopus, we chose two genes for each specie: z*Urod*, z*Tyr* for zebrafish and x*P48*, x*Tyr* for xenopus. In addition, four target sites of each gene were designed. Validated CRISPR/Cas9 targets were used as positive control(Guo et al., 2014; Qin et al., 2015). The data of mRNA injected embryos show no significant occurrence of DSB at these target sites (**Supplementary Figure 3a, b, c, d**). There is a suspected difference between control groups and one experimental group (**Supplementary Figure 3a**). After Sanger sequencing, those are polymorphism at non-target area (data no to show). So, we suspected this result may be because that the number set of targets we detect is too small to be representative of AsCpf1 capacity. So we extended the target sets of these two species, the results show that there is still no DSB present (**Supplementary Figure 3e, f, g, h**).

In order to understand whether the target selection or some other issue is the reason of these negative results. We designed an exogenous target detection(ETD) system: cloned the mouse *Tet2* Cpf1 target into the vector that expresses GFP, named pTg-mTet2 (**Supplementary Figure 4a**), it was co-transfected with AsCpf1 expression plasmid and crRNA into human HEK293T cells. The results showed that exogenous m*Tet2* target was edited in human cells (**Fig d**, **Supplementary Figure 4b**). After verifying the ETD system can effectively detect the effectiveness of exogenous target, we chose several zebrafish and xenopus targets to test on this system. Surprisingly, it has been found that all those targets are high cleavage efficiency in the HEK293T cells (**Fig d**, **Supplementary Figure 4d**).

Exogenous target detection experiments have shown that the appearance of negative results is not due to target selection. So we turned our attention to other possible reasons. The culture temperatures of zebrafish and x enopus were less than 30 °C, which is far below the culture temperature of the two mammalian embryos. This makes us wonder if temperature is the deciding factor - AsCpf1 protein activity was lacked in the non-optimal working environment. So we did the in vitro enzyme activity test. A certain unit of purified AsCpf1 protein can be very effective in cutting DNA at 37 °C (**Supplementary Figure 5a, b**). The same goes for reactions that add embryonic lysates at 37 °C, however, our experimental results showed that the activity of AsCpf1 was significantly lower at 28.5 °C and 25 °C (**Fig e**).

Since injected mRNAs take a few hours to translate into protein in the embryos, and AsCpf1 protein is greatly reduced in activity at low temperatures, the strategy of injecting mRNAs is apparently inappropriate. We used crRNAs from those target that had been detected in the ETD system, mixed with AsCpf1 protein and injected into xenopus and zebrafish embryos. By T7E1 DSB assay, there was a slight difference between the experimental and control groups in each of the two species (**Fig f, g**). We confirmed that there was a single-base mutation in both experimental groups by sanger sequencing (**Fig h, i**).

One finer temperature gradient experiment showed that AsCpf1 protein had the highest activity at about 37 °C(**Fig j**, **Supplementary Table 5, 6**), while had a significant activity decrease as the temperature increased or decreased (**Fig k**, **Supplementary Table 7**). We believe that temperature may be the reason why AsCpf1 cut only a small amount of DNA in the two low temperature species-temperature sensitivity.

CRISPR/Cpf1 system has significant advantages in working mechanism. Our data reveal that AsCpf1/crNA system is effective at some target sites in the mammalian embryos. Unfortunately, it did show only a very low-level function in zebrafish and xenopus, which are two classic developmental biology animal models. This may be due to the sensitivity of AsCpf1 to temperature. And prolonged the reaction time under low temperature conditions is an effective way (**Supplementary Figure 5c**), which can also explain why the low temperature species that have been reported can effectively be genetically modified after obtaining transgenic lines(Port and Bullock, 2016; Tang et al., 2017). But, it is still unknown that whether to extend the reaction time in non-optimal conditions will increase the risk of off-target? If scientists want to develop CRISPR/Cpf1 system into a mature gene-editing tool, more test and optimization on different species are needed. In this study, we only discussed the effectiveness of the CRISPR/Cpf1 systems in four different animal models, and did not touch the problem of off-target. As a critical evaluation criteria of genome editing tool, off-target of CRISPR/Cpf1 system remains to be further studied.

## METHODS

### RNA in vitro transcription

AsCpf1 was synthesized by IGE Biotech Ltd. and cloned into the pCS2+ vector. AsCpf1 and SpCas9 mRNA were transcribed using the mMessage mMachine SP6 Kit (Thermo Fisher). All crRNAs and sgRNAs were synthesized using the TranscriptAid T7 High Yield Transcription Kit (Thermo Fisher). ssDNA oligos (Supplemental Material) corresponding to the reverse complement of the target sequence were synthesized from Thermo Fisher Scientific Inc and IDT, and annealed to a short T7 primer. All RNAs were purified using the ZR-96 RNA Clean Kit (Zymo).

### mESCs culture and transfection

Mouse embryonic stem cells (mESCs) were maintained on feeder layers or feeder free with mESC-2i medium (DMEM, 15%FBS, NEAA, GlutaMAX, PD0325901, Chir99021, LIF) at 37°C with 5% CO_2_. Plasmids were transfected by Mouse ES Cell Nucleofector kit (Lonza) following the manufacturer’s instruction. RNAs were transfected by Lipofectamine RNAiMAX (Thermo Fisher) following the manufacturer’s instruction, and ribo*TRACER*™ Green (Ribobio) as the positive control.

### HEK293T cells culture and transfection

HEK293T cell lines were maintained in Dulbecco’s modified Eagle’s medium (DMEM) supplemented with 10% FBS (HyClone) at 37 °C with 5% CO_2_ incubation. Plasmids were transfected by Lipofectamine 3000 (Thermo Fisher) following the manufacturer’s instruction. RNAs were transfected by Lipofectamine RNAiMAX (Thermo Fisher) following the manufacturer’s instruction.

### Mouse embryos acquisition, RNAs microinjection, and culture

All animal experiments and Protocols were approved by the Institutional Animal Care and Use Committees (IACUC) of the Laboratory Animal Research Center at Guangzhou Institutes of Biomedicine and Health (Permit Number:2015009). C57BL/6 (B6) mouse strains were used as embryo donors. Female B6 mice (6-8 weeks old) were super-ovulated by intraperitoneal injections of 5 IU pregnant mare serum gonadotropin (PMSG, Ningbo Renjian Pharmaceutical Co., Ltd, Ningbo, China) and 5 IU human chorionic gonadotropin (hCG, Ningbo Renjian Pharmaceutical Co., Ltd, Ningbo, China) at 48-hour intervals. The fertilized embryos were collected from oviducts of super-ovulated female after mating with B6 stud males. RNAs (100ng/uL Cpf1/Cas9 mRNA and 50ng/uL crRNA/sgRNA) was injected into the cytoplasm of fertilized eggs with well recognized pronuclei in M2 medium (Millipore). The injected zygotes were cultured in G1/G2 medium (Vitrolife) at 37°C under 6% CO_2_ in air until blastocyst stage by 3.5 days.

### Pig oocyte/embryos acquisition, activation, RNAs microinjection, and culture

Briefly, pig ovaries were collected from a local slaughterhouse and transported to the laboratory in 0.9% saline with penicillin and streptomycin at 35 °C to 39 °C. Cumulus oocyte complexes(COCs) were aspirated from antral follicles using 10ml syringe. After washed 3 times using maturation medium, Tissue Culture Medium 199 (Gibco), which was supplemented with 0.1% (w/v) polyvinyl alcohol (Sigma), 3.05 mM D-glucose, 0.91 mM sodium pyruvate (Sigma), 0.57 mM cysteine (Sigma), 0.5 mg/mL luteinizing hormone (LH Sigma), 0.5 mg/mL follicle stimulating hormone (FSH Sigma), 10 ng/mL epidermal growth factor (Sigma), 10% (v/v) porcine follicular fluid, 75 mg/mL penicillin G, and 50 mg/mL streptomycin, the COCs were transferred to maturation medium for 42-44 h. Then, the matured oocytes were released from the cumulus cells by vigorous vortexing for 5 min in TL-HEPES containing 0.1% hyaluronidase (Sigma) and actived in activation medium (0.3 M mannitol, 1 mM CaCl_2_•2H_2_O, 0.1 mM MgCl_2_•6H_2_O, and 0.5 mM HEPES) with 2 successive DC pulses of 120 V/mm for 30 μseconds using an electrofusion instrument (CF-150B, BLS, Hungary). After recovered for 30 mins, the active oocytes were injected with 200ng/uL Cpf1/Cas9 mRNA and 20ng/uL crRNA/sgRNA and ddH_2_O as control. The injected embryos were cultured in PZM3 for 6 days and the blastocysts were collected for analysis.

### Zebrafish embryos microinjection and culture

Wild type Tu line embryos were raised at 28.5°C. One-cell stage zebrafish embryos were injected with 2 nL of a solution containing 200 ng/μL Cpf1/Cas9 mRNA/protein and 20 ng/μL crRNA/sgRNA. For exogenous targets test, injection solution was included 10 pg/uL pTg-mTet2 plasmid. The injected embryos were cultured in 1× E3 mediumv (5.0 mM NaCl, 0.17mM KCl, 0.33mM CaCl, 0.33mM MgSO_4_, pH 7.4) for 2 days, and then were collected for analysis.

### Xenopus embryos microinjection and culture

One-cell stage xenopus embryos were injected with 2 nL of a solution containing 300ng/μL Cpf1/Cas9 mRNA/protein and 100ng/μL crRNA/sgRNA. For exogenous targets test, injection solution was included 10 pg/uL pTg-mTet2 plasmid. The injected embryos were cultured in 0.1× MBS medium (1× MAS: 88 mM NaCl, 2.4 mM NaHCO_3_, 1 mM KCl, 0.82mM MgSO_4_, 0.33 mM Ca(NO_3_)_2_, 0.41 mM CaCl_2_, 10 mM HEPES pH 7.4) for 24 hours, and then were collected for analysis.

### T7E1 DSB assay

Genomic DNA was extracted using the QuickExtract DNA Extraction Solution (Epicentre) following the manufacturer’s instruction. Genomic region surrounding the target sites for each gene was PCR amplified. Half PCR products were used for T7 Endonuclease I assay: 10uL PCR products were mixed with 2uL NEB buffer 2 and ddH20 to a final volume of 20uL, and subjected to a re-annealing process to enable heteroduplex formation: 95°C for 10min, 95°C to 85°C ramping at - 2°C/s, 85°C to 25°C at −0.2°C/s, and 25°C hold for 1 minute. After re-annealing, products were treated with T7 Endonuclease I (NEB) at 37 °C for 30min, and analyzed on 10% TBE poly-acrylamide gels. Gels were stained with SYBR Gold DNA stain (Thermo Fisher) for 30 minutes and imaged with Tanon 1600 digital gel image analysis system (Tanon). The remaining PCR products were gel extraction, cloned into pMD18T (Takara) vector, and sequenced for counting indels.

### In Vitro Enzyme Activity Assay

Cpf1 (with Flag tag) expression plasmid was transfected into HEK293T cells as described below. Cytoplasmic lysates from HEK293T cells were prepared with NE-PER Nuclear and Cytoplasmic Extraction Reagents (Thermo Fisher) supplemented with Protease and Phosphatase Inhibitor (Thermo Fisher). Flag tag protein was purified by FLAG M purification mammalian expression systems (Sigma-aldrich) following the manufacturer’s instruction. The in vitro enzyme activity assay was carried out as follows: for a 20 uL cleavage reaction, a certain volume of endonuclease protein was incubated with 2uL 10X FastDigest Buffer (Thermo Fisher), 0.5 ug in vitro transcribed RNA and 250ng SalI-linearized pTg-mTet2 plasmids. After incubation, cleavage reactions were stoped by 6X Reaction Stop Solution (60 mM EDTA, 20 mM pH 8.0 Tris-HCl, 0.48% SDS), purified using ZR-96 DNA Clean & Concentrator (Zymo Research) and analysed on 10% TBE poly-acrylamide gels.

## Acknowledgements

This work were supported by National Natural Science Foundation of China (31421004, 91419310, 31530038, 31471367, 31271554), National Basic Research Program of China (973 Program of China, 2012CB966802, 2015CB942803), Guangdong Science and Technology Project (2014B050504008, 2014B050502012, 2014B030301058, 2014A030304010). The Science and Technology Planning Project of Guangdong Province (2016A030303046), and Pearl River Science and Technology Nova Program of Guangzhou (201710010112).

